# Differential regulation and production of secondary metabolites among isolates of the fungal wheat pathogen *Zymoseptoria tritici*

**DOI:** 10.1101/2021.08.12.456184

**Authors:** M. Amine Hassani, Ernest Oppong-Danquah, Alice Feurtey, Deniz Tasdemir, Eva H. Stukenbrock

## Abstract

The genome of the wheat pathogenic fungus, *Zymoseptoria tritici,* represents extensive presence-absence variation in gene content. Here, we addressed variation in biosynthetic gene clusters (BGCs) content and biochemical profiles among three isolates. We analysed secondary metabolite properties based on genome, transcriptome and metabolome data. The isolates represent highly distinct genome architecture, but harbor similar repertoire of BGCs. Expression profiles for most BGCs show comparable patterns of regulation among the isolates, suggesting a conserved “biochemical infection program”. For all three isolates, we observed a strong up-regulation of an abscisic acid (ABA) gene cluster during biotrophic host colonization, indicating that *Z. tritici* potentially interfere with host defenses by the biosynthesis of this phytohormone. Further, during *in vitro* growth the isolates show similar metabolomes congruent with the predicted BGC content. We assessed if secondary metabolite production is regulated by histone methylation using a mutant impaired in formation of facultative heterochromatin (H3K27me3). In contrast to other ascomycete fungi, chromatin modifications play a less prominent role in regulation of secondary metabolites. In summary, we show that *Z. tritici* has a conserved program of secondary metabolite production contrasting the immense variation in effector expression, some of these metabolites might play a key role during host colonization.

**Originality-Significance Statement:** *Zymoseptoria tritici* is one of the most devastating pathogens of wheat. So far the molecular determinants of virulence and their regulation are poorly understood. Previous studies have focused on proteinasous virulence factors and their extensive diversity. In this study, we focus on secondary metabolites produced by *Z. tritici*. Using a comparative framework, we here characterize core and non-core metabolites produced by *Z. tritici* by combining genome, transcriptome and metabolome datasets. Our findings indicate highly conserved biochemical profiles contrasting genetic and phenotypic diversity of the field isolates investigated here. This discovery has relevance for future crop protection strategies.

## Introduction

Fungal genomics has revealed a large and untapped diversity of biosynthetic gene clusters (BGCs) encoding secondary metabolites (Rokas *et al*., 2020). These metabolites have a broad spectrum of biological functions and play an essential role as determinants of fungal life style and niche specialization. Most of the secondary metabolites produced by fungi derive from non-ribosomal peptides, polyketides or mixed polyketide–non-ribosomal peptides or other biosynthesis pathways (such as terpenes) (Brakhage, 2013). Genes involved in the biosynthesis of secondary metabolites are often physically linked in gene clusters allowing co-regulation of gene expression and metabolite production (Rokas *et al*., 2018). Commonly, BGCs encode for the enzymes that are responsible for biosynthesis of the metabolite backbone including nonribosomal peptide synthetases (NRPS), polyketide synthases (PKS), and fusions of PKS and NRPS (PKS-NRPS/NRPS-PKS hybrids), as well as other enzymes involved in further modifications of the metabolite backbone. Moreover, certain BGCs encode genes involved in the metabolite transport and/or genes conferring resistance to the activity of the metabolite.

Plant-associated fungi produce a multitude of secondary metabolites to facilitate host invasion, to manipulate host defenses, interact with microorganisms and to sequester essential nutrients in plant tissues (Scharf *et al*., 2014). Moreover, many fungi, including plant pathogens, produce plant hormones such as gibberellins, abscisic acid (ABA), auxin and cytokinin, which potentially may interfere with the hormone balance of plants that are colonized by these fungi. However the regulation and relevance of these secondary metabolites in plant-fungus interactions is still little known (Reineke *et al*., 2008; Chanclud *et al*., 2016; Tudzynski and Hölter, 1998).

Genome data has provided detailed insights into the diversity in BGCs among closely related plant-associated fungi (e.g. (Schardl *et al*., 2013; Wight *et al*., 2013; Villani *et al*., 2019)), however to which extent, and how, this diversity is translated into chemical diversity has only been addressed in a few species. Genome sequencing across diverse taxa has shown that the organization of BGCs in fungal genomes can be highly variable between closely related species and in some cases even between different strains of the same species. Members of the fungal family Clavicipitaceae includes plant-associated species with diverse life styles ranging from mutualistic symbionts and endophytes to pathogens. This group of fungi exhibits an exceptional chemotypic diversity originating from diverse architecture of gene clusters responsible for secondary metabolite production (Schardl *et al*., 2013). Comparative genome analyses in other groups of ascomycete plant pathogens have likewise revealed structural variation associated with BGCs, for example in the gene clusters responsible for synthesis of the phytotoxins botrydial and botcinic acid in *Botrytis* species (Valero-Jiménez *et al*., 2020) and in gene clusters responsible for the synthesis of distinct secondary metabolites, including the mycotoxin fucosarin, in isolates of *Fusarium fujikuroi* (Niehaus *et al*., 2016). Possibly this variation in BGC content and composition reflect the effect of diversifying selection acting on genes involved in antagonistic plant pathogen interactions.

Fungi accommodate different environmental and ecological conditions by regulating and optimizing the production of secondary metabolites. This regulation can occur at different levels from individual genes and clusters to global regulation of BGCs and involve complex networks of regulatory components (Brakhage, 2013). In several fungal pathogen taxa including species of *Fusarium* and *Epichloë* chromatin modifications such as histone methylation or acetylation have been shown to play an important role as global regulators of BGCs as well as regulators of specific gene clusters (Connolly *et al*., 2013; Studt *et al*., 2013; Chujo and Scott, 2014; Lukito *et al*., 2019). In the fungal pathogen *Fusarium graminearum*, secondary metabolism is largely regulated by the silencing histone mark H3K27me3. This was demonstrated by deletion of the histone methyltransferase *kmt6* responsible for the heterochromatin-associated tri-methylation of lysine 27 on histone H3 (H3K27me3). The mutant exhibited a constitutive expression of secondary metabolite cluster along the genome severely impacting fungal fitness (Connolly *et al*., 2013). The cluster organization of biosynthesis genes has been proposed to favor a chromatin based regulation of secondary metabolite production as it allows transcriptional activation in narrowly restricted regions (Brakhage, 2013).

We have recently described extensive variation in genome composition among closely related species of grass pathogens in the genus *Zymoseptoria* (Feurtey *et al*., 2020). This genus includes the important wheat pathogen *Z. tritici* that has devastating impact on wheat production world-wide. Population genomic sequencing of *Z. tritici* has revealed extensive genetic variation along the genome (Plissonneau *et al*., 2018), and it is hypothesized that this variation allows the fungus to rapidly adapt to changes in the environment including new host resistances and fungicide treatments applied in wheat fields. *In silico* predictions of gene clusters have demonstrated how BGCs are enriched in the more rapidly evolving sub- telomeric regions of the genome of *Z. tritici,* which may facilitate genetic diversification of the BGCs (Cairns and Meyer, 2017). *Z. tritici* is also characterized by an exceptional high variability in growth morphology, for example some isolates being highly melanized and others non-melanized when grown under laboratory conditions (Fagundes *et al*., 2020). The extensive genetic and phenotypic variability manifested by this fungus led us to hypothesize a comparably high variation in the overall secondary metabolite production.

In this study, we explore genomic diversity of three isolates of *Z. tritici* to address the composition of BGC along the fungal genome. We complement genomic analyses with transcriptome and metabolome data to compare potential variations in secondary metabolite profiles of the three field isolates. Interestingly, we find evidence for conserved biochemical profile of the three isolates contrasting the highly variable content and production of effector proteins. Lastly, we analyse secondary metabolite production in a *kmt6* mutant (deletion of the KMT6 histone methyltransferase gene) impaired in catalyzing the histone modification H3K27me3 to assess the relevance of chromatin-based BGC regulation in *Z. tritici*. We provide evidence for extensive chemical differences between the three field isolates of *Z. tritici*, and show that histone modifications in this fungus, in contrast to other fungi such as species of *Fusarium* and *Aspergillus* (Keats *et al*., 2007; Smith *et al*., 2011; Studt *et al*., 2013; Pidroni *et al*., 2018), plays a minor role in regulating secondary metabolite production during axenic growth.

## Results

### *In silico* predictions and comparison of BGCs among three field isolates of *Z. tritici*

To compare the genomic potential for secondary metabolite production among isolates of *Z. tritici*, we used the program antiSMASH (Blin *et al*., 2017) to predict the distribution of BGCs in high quality genome assemblies of three field isolates Zt05, IPO323 and Zt10 (Feurtey *et al*., 2020). We have previously described and compared the infection development of Zt05, IPO323 and Zt10 in susceptible wheat and demonstrated that these isolates, although morphologically and genetically highly distinct, are equally virulent in a susceptible wheat cultivar (Haueisen *et al*., 2018). Interestingly, these three isolates exhibit an exceptional phenotypic diversity when cultivated on agar whereby the isolate Zt10 appears highly melanized in contrast to the two other isolates (Figure 1A).

**Figure 1.**
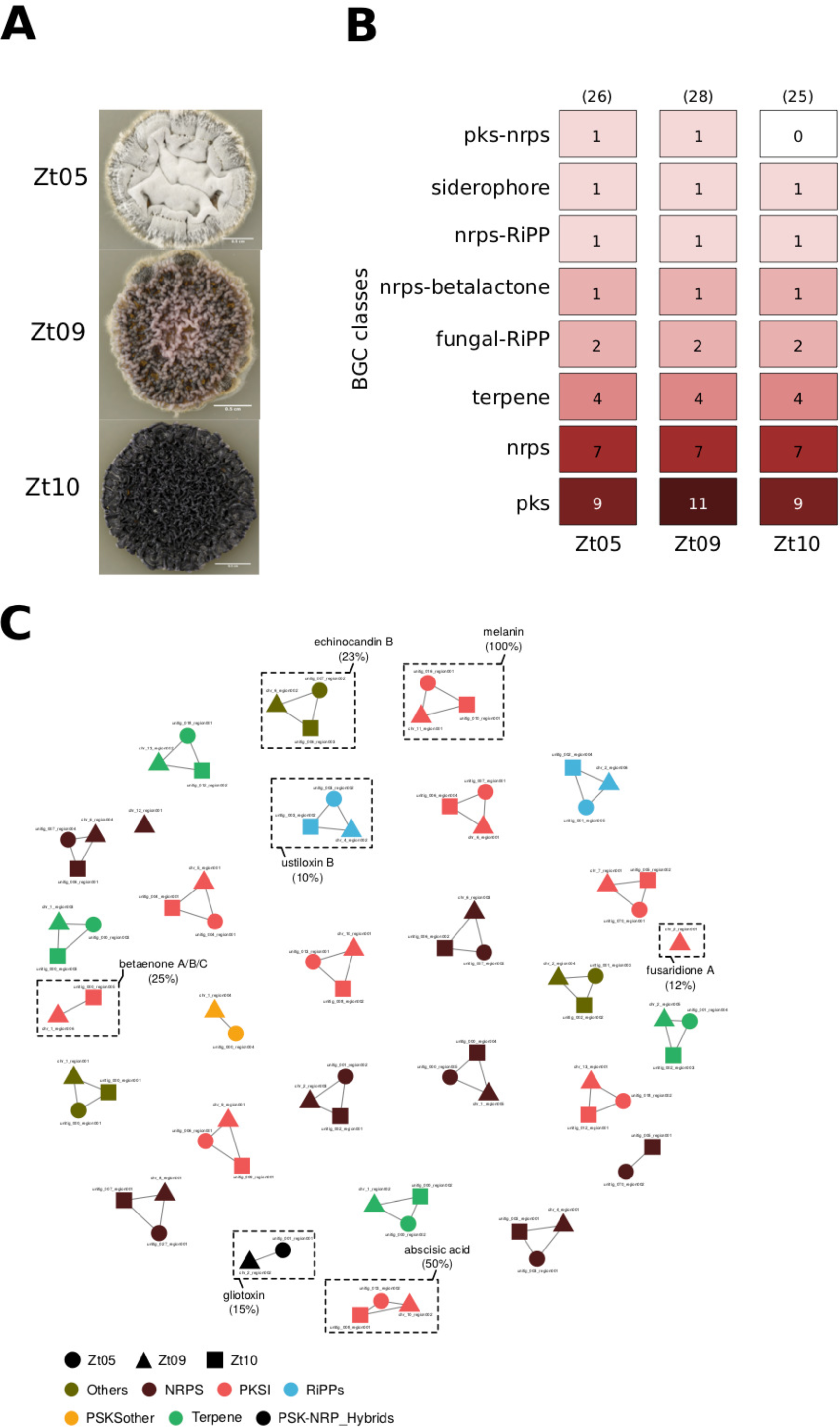
Mining the genome of three *Z. tritici* field isolates for biosynthetic gene clusters. **A)** Differences in the colony morphology between the three *Z. tritici* isolates (i.e., Zt05, IPO323 and Zt10) after 15 days of growth on solid medium (YMS) at 18° C. **B)** Output from antiSMASH analysis that predicts biosynthetic gene clusters (BGCs) from the genome of the three *Z. tritici* isolates. The fungal isolates Zt05, IPO323, and Zt10 are predicted to harbor 26, 28, and 25 BGCs, respectively. These BGCs are categorized into eight classes including Polyketide synthase (pks), Non-ribosomal peptide synthetase (nrps), terpene, ribosomally synthesized and post-translationally modified peptides (fungal-RiPPs), siderophore and hybrid nrps-betalactone, nrps-RiPP and pks-nrps clusters. **C)** Similarity network of BGCs predicted from Zt05 (circle), IPO323 (triangle) and Zt10 (square). Each node in the network (i.e. circle, square or triangle) depicts a BGC and the color indicates the class. BGCs connected by edges belong to a same biosynthetic gene family. Seven BGFs (highlighted with dashed-rectangle) showed percent of gene similarity to annotated clusters in the MiBIG databese.

Based on the *in-silico* prediction, we find that the three isolates overall encode comparable numbers of BGCs (Figure 1B). In total, we identified 26, 28 and 25 BGCs in the genomes of Zt05, IPO323 and Zt10, respectively. These gene clusters are categorized into several classes including polyketide synthases (PKS), non-ribosomal peptide synthetases (NRPSs), terpene synthase (TPS), fungal RiPPs (ribosomally synthesized and post-translationally modified peptides), hybrid NRPS-betalactone, NRPS-RiPPs and PKS-NRPSs and siderophores (Figure 1B, Table S1, Table S2 and Table S3).

We next investigated in details the variation in BGC distribution and sequence composition among genomes of the three isolates. To this end, we calculated the similarity across all predicted BGCs using BiG-SCAPE to further identify shared and unique BGCs within and among the *Z. tritici* isolates (Navarro-Muñoz *et al*., 2020). The resulting sequence similarity network consists of 29 biosynthetic “gene families” (BGFs), shown as sub-network (Figure 1C) and representing seven main BGC classes PKS, NRPS, RiPPs, PKS-NRP hybrids, TPS, “other PSKs” and “others”. Almost all BGFs (27) count two to three BGCs, and none of these BGFs are composed of two BGCs originating from the same isolate (Table S4). We identified two BGCs in IPO323 that show no similarity to other BGCs (using a clustering cutoff of 0.3), indicating that this isolate exhibits to some extent a unique biosynthetic potential. However, the overall *in silico* BGC predictions suggest that the three isolates share overall similar biochemical potential.

Of all predicted BGCs, we observed that five, seven and five BGCs, in Zt05, IPO323 and Zt10, respectively, show similarity to annotated BGCs in the MiBIG database (with a similarity > 10% in terms of gene content and order) (Figure 1C, Table S1, Table S2 and Table S3, respectively). These annotated gene clusters have similarity to gene clusters producing the compounds betaenone, ustiloxin, fusaridione, gliotoxin, echinocandin, melanin and absicic acid. Several of these compounds are likely to play a role in pathogenicity of *Z. tritici*: betaenone and ustiloxin have previously been described as phytotoxins in the plant pathogenic fungi *Phoma betae* and *Ustilaginoidea virens*, respectively (Ichihara *et al*., 1983; Brauers *et al*., 2000; Abbas *et al*., 2014). Fusaridione and gliotoxin, mainly described in the plant pathogenic fungus *Fusarium heterosporum* and the human pathogen *Aspergillus fumigatus*, respectively, are known to have antimicrobial properties that may facilitate competitiveness of *Z. tritici* with respect to other plant-associated microbes (Kakule *et al*., 2013; Svahn *et al*., 2014). Echinocandins are fungal-produced metabolites that are responsible for antifungal activity against β-(1,3)-D-glucan synthesis (Walker *et al*., 2010). Finally, melanin is known to be a critical component of pathogenicity in many plant pathogenic fungi (Nosanchuk and Casadevall, 2003). Our *in silico* prediction moreover identified one BGC present in all three isolates which shows 50% similarity to an abscisic acid (ABA) gene cluster described in *Botrytis cinerea* (Table S1, Table S2 and Table S3) (Izquierdo Bueno *et al*., 2018). Abscisic acid is also known as a plant hormone involved in abiotic and biotic stress regulation, and this hormone can act to suppress defense regulation (e.g. (Mohr and Cahill, 2003; Anderson *et al*., 2004; Koga *et al*., 2004). We speculate that it may be produced by *Z. tritici* to interfere with defense regulation in wheat during pathogen infection.

### Differential regulation of Biosynthetic Gene Clusters (BGCs) in three field isolates of *Z. tritici*

We next asked how secondary metabolite production is regulated during plant infection in the three *Z. tritici* isolates. To this end, we used previously published transcriptome data generated for the three isolates during wheat infection (Haueisen *et al*., 2018). The infection development of *Z. tritici* is characterized by four infection stages previously described with detailed microscopy analyses (Haueisen *et al*., 2018). In brief, stage A represents the initial penetration of the fungus via the stomata and early colonization of the mesophyll tissue. Stage B represents the biotrophic colonization of the mesophyl tissue, stage C represents the transition from biotrophic to necrotrophic growth; and finally stage D represents the necrotrophic colonization and asexual reproduction.

We focused our gene expression analyses on the BGCs identified by our *in-silico* prediction. To this end, we first show that the vast majority of predicted BGCs are expressed during plant colonization (between 87% of genes in stage A for Zt10 and 98% in stage D for Zt09 (a clone derived from IPO323), Figure S1). Consistent with the low amount of fungal biomass and therefore overall lower proportion of fungal transcripts, we found an increased transcript abundance of all genes, including BGC genes in the later stages of infection (C and D) (see percentage of alignments increasing in Table S5). Next, we compared the expression specifically of BGC genes in the three isolates throughout the infection development to uncover patterns of secondary metabolite production (hereby we compared the differential expression between the consecutive stage A and B but not between stage A and D). This comparison revealed a dynamic expression of BGC genes with 9%, 13% and 8% of the BGC genes in Zt05, Zt09 and Zt10, respectively being differentially expressed between the infection stages (Figure 2A, Table S6). In comparison, the overall proportion of differentially expressed genes for non-BGCs were consistently lower, 2.6%, 4.5% and 2.3% for Zt05, Zt09 and Zt10, respectively, suggesting that different BGCs are important during different infection stages of *Z. tritici*.

**Figure 2.**
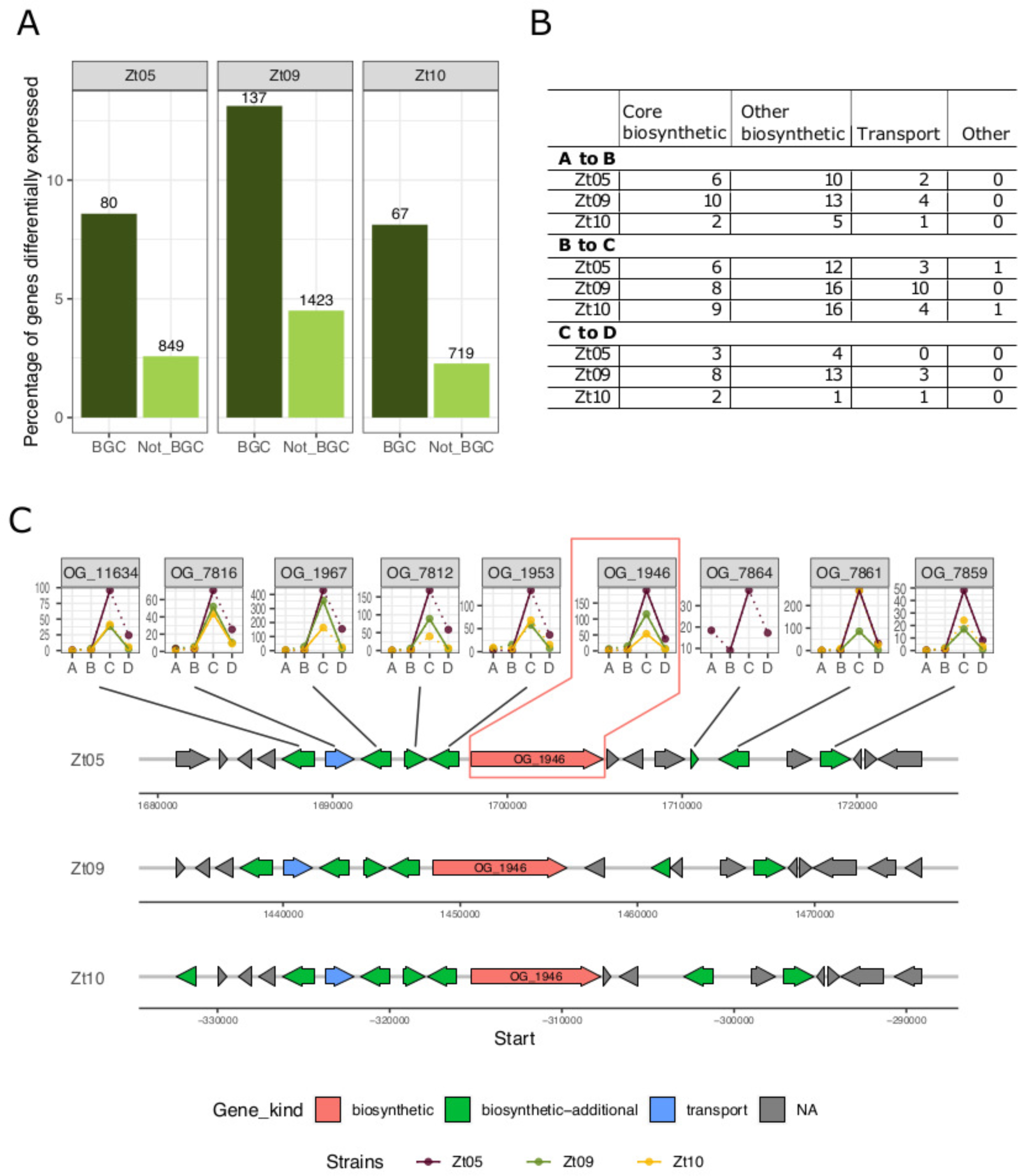
Expression profiles of genes associated with biosynthetic gene clusters in *Z. tritici* isolates. **A)** Bar plot representing the percentage of genes differentially expressed during host infection (based on comparison of four infection stages). The numbers above the bars represent the number of differentially expressed genes predicted to be in gene clusters (“BGC” in dark green) and all other genes (“Not BGC” in light green). **B)** Numbers of differentially expressed genes in BGCs for each *Z. tritici* isolate during four infection stages. **C)** Schematic overview of the genetic architecture of the BGC predicted to produce abscisic acid (ABA). The expression kinetics of the ABA genes in the three isolates are represented as line plots, linked to the orthologs in Zt05. On the line graphs, the y-axis represents the TPM values while the x-axis denotes the infection stages. A dotted line represents change in expression level that is not significant whereas significant differences are represented by a full line. The line plots are labeled with the orthogroup number as identified in (Feurtey *et al*., 2020).

The genes encoded by individual BGC have different functions in the metabolite syntheses, and we next focused on the expression of individual genes in the BGCs predicted to have a primary function in the metabolite synthesis. The program antiSMASH distinguishes: core biosynthetic genes, biosynthetic genes, regulatory genes, transporters, and genes with unknown function for individual BGCs. We focused specifically on the core biosynthetic genes from the predicted BGC using previously defined orthogroups (Feurtey *et al*., 2020) to compare expression patterns between isolates (Figure 2B). In total, we identified 44 core biosynthetic gene orthogroups among the 29 BGCs (found in at least one strain). Of note, a few clusters contain more than one core biosynthetic gene and some clusters were identified in only one or two of the isolates. We investigated the expression kinetics of these genes throughout the four infection stages and revealed a highly dynamic expression pattern of the core BGC genes (Figure S2, Table S7). In particular, we observed a general pattern of higher gene transcription during the stages C or D, i.e., the necrotrophic phase, an increase in transcription that is significant in several core biosynthetic genes (see for example, BGFs 8, 9, 15, 18, 20, 24 in Figure S2).

We also identified differences in expression of BGC genes among the three isolates. For example, one gene in the BGC encoding the phytotoxin betaenone (BGF5:OG_10647) show an increased expression in the isolates Zt10 during the transition from biotrophic to necrotrophic host colonization suggesting different relevance of the metabolite among the three *Z. tritici* isolates. We compared the expression of seven genes annotated in the gene cluster BGF_11 responsible for the biosynthesis of ustiloxin. Overall, the genes were expressed in the same manner in the three isolates. Six of the seven ustiloxin biosynthesis genes showed an increased expression during stage B and C suggesting that this phytotoxin may play a role in the transition from biotrophic to necrotrophic host colonization. One gene BGF 11:OG_11282 on the other hand show a low expression during stages A to C, but then a strong up-regulation during the late phase of infection and necrotrophic host colonization. Genes annotated in the BGC involved in the biosynthesis of the two antimicrobial compounds fusaridione (BGF_28) and gliotoxin (BGF_29) did not show a consistent expression pattern during host colonization of the three isolates. The gene BGF_28:OG_5759 putatively involved in fusaridione biosynthesis showed a decreased expression from stage A through D in Zt09. The gene BGF_29:OG_2829 showed an increased expression in Zt05, but a more dynamic and inconsistent pattern in Zt09. Three of four genes annotated in the BGC predictive to confer the biosynthesis of echinocandin (BGF17) showed an up-regulation during early biotrophic host colonization in the three isolates while one gene BGF_17:OG_5907 on the other hand was down- regulated (in the isolate Zt09 only). One gene predicted to be involved in melanin biosynthesis and encoded by the BGC_23 showed an increased expression in all three isolates throughout the infection development (stages A to D), possibly reflecting protection of the fungal cell wall during infection establishment and progress.

One gene showing a strong up-regulation in the three isolates during the transition from biotrophic to necrotrophic growth (i.e., phase C) belongs to the BGF_20 predicted to be responsible for the production of ABA (BGC showing 50% gene similarity to the abscisic acid gene cluster in *Botrytis cinere*a). Given the potential role of ABA in pathogenicity, we further investigated the genetic architecture of the ABA BGC in *Z. tritici* and transcription among the three isolates, as well as the prediction function of each gene (Figure 2C, Table S8). Overall, we found that the synteny and gene functions of the predicted ABA BGC are conserved among the three isolates: eight genes with a predicted role in the cluster function were shared in the three isolates and have the same order, however one additional gene was found in the BGC of Zt05 and another in Zt10 (Figure 2C). The expression kinetics of the genes in the ABA cluster predicted to have a function in the compound synthesis and in transport showed a common up-regulation at the onset of the necrotrophic phase (significant up-regulation in two isolates for all shared orthologs, and significant in all three isolates for the core biosynthetic gene) (Figure 2C, Tables S5 and S6).

The expression patterns of the ABA genes, which we observe *in planta,* suggest that ABA is produced by *Z. tritici* during host infection to manipulate plant defenses. We furthermore hypothesize that the other BGC genes, which show a similar kinetics, likewise play an important role during host colonization.

### Variation in secondary metabolite profiles of *Z. tritici* field isolates

We next sought to characterize the chemical profile of the three field isolates during axenic growth. To this end, we applied a comparative untargeted metabolomics approach using MS/MS (MS^2^)-based molecular networking (MN). MN is a comprehensive algorithm-based metabolomics tool that compares MS^2^ spectra and categorizes them into different structural clusters based on fragmentation patterns (Wang *et al*., 2016). The recently developed tool, feature based molecular networks (FBMN), incorporates MS^1^ related information such as retention time and isotope patterns into data processing, thereby facilitating the annotation of molecular families (MF) and allowing for resolution of positional isomers and relative quantification (Soyer *et al*., 2014; Nothias *et al*., 2020; Wang et al, 2016). Herein we applied FBMN analyses that assisted us to determine and compare the secondary metabolite profile of the three *Z. tritici* field isolates.

The metabolome analyses of the *Z. tritici* field isolates revealed a total of 170 the so-called nodes, each representing one molecular ion in the crude metabolite extracts (see Materials and Methods). We grouped these into 21 clusters and 43 singletons (Figure 3A). Overall, five putative metabolite clusters were annotated as polyketide (PK), fatty acid (FA), nonribosomal peptide (NRP) and terpene (TER) MFs, as also identified in the genome analyses. The data from MN was used to generate a Venn diagram to visualize the metabolite distribution among the isolates (Figure 3B). We find that Zt05 displayed 138 ions, IPO323 132 ions and Zt10 152 ions. Twenty-one ions were exclusively found in one isolate, namely ten ions for Zt10, six ions for Zt05 and five ions for IPO323 respectively (Figure 3B and Table S9). These strain-specific ions could be responsible for individual strain variation in biochemical profiles, for example the very prominent variation in melanization (Figure 1A). Conversely, a total of 103 ions were common to all field isolates while 46 ions were shared by two strains (Figure 3B). These data indicate that the field isolates overall produce similar metabolites when maintained under the same culture conditions.

**Figure 3.**
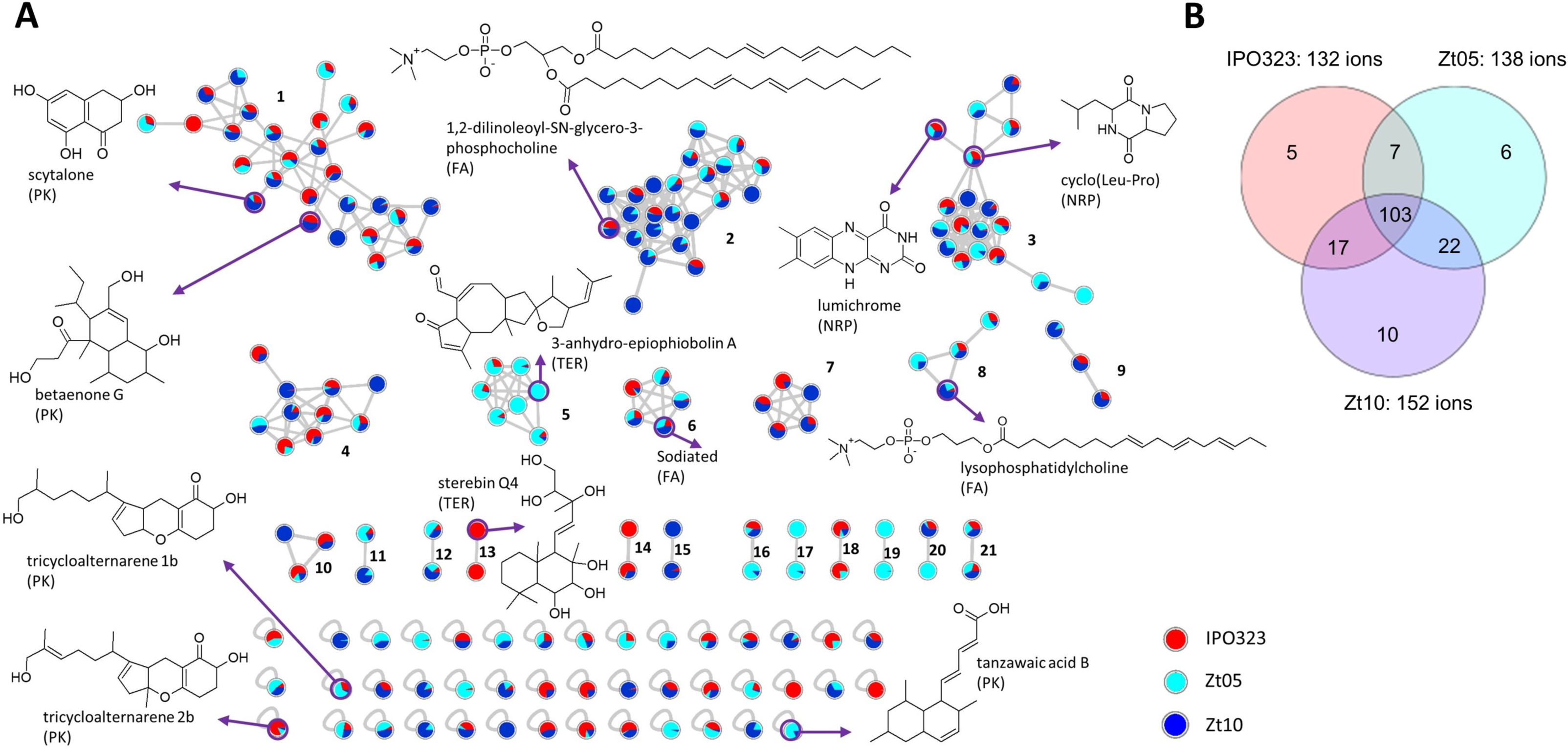
Comparative metabolomics of three *Z. tritici* field isolates. **A)** Molecular network (MN) generated from the Global Natural Product Social MN platform of crude organic extracts from the three isolates of *Z. tritici*. Nodes represent molecular ions detected in the crude extracts with colour coding indicative of the relative abundance in each *Z. tritici* sp. (red-IPO323, light blue-Zt05 and deep blue-Zt10. Some annotated compounds are displayed with their chemical classes: polyketide (PK), nonribosomal peptide (NRP), terpene (TER) and fatty acid (FA). Other annotations are displayed in Table S8. **B)** Venn diagram displaying specific and shared parent ions detected in the culture extracts of the three *Z. tritici* spp.

As shown in Figure 3A, a combined automated and manual dereplication effort resulted in the putative annotation of two polyketides, scytalone (Nakamura *et al*., 2020) and betaenone G (Schlörke, 2005), a diketopiperazine (NRP) cyclo Leu-Pro (Tunes et al., 2019), the labdane diterpene sterebin Q4 (Kamauchi et al., 2015), phospholipids lysophosphatidylcholine (Vijayakumar et al., 2016) and 1,2-dilinoleoyl-SN-glycero-3-phosphocholine (Mills and Goldhaber, 2010). Other annotated compounds included three PKs, the benzopyranones trycycloalternarene 1b and tricycloalternarene 2b (Sun *et al*., 2013), the bicyclic tanzawaic acid (Matsuo, et al. 2020) and the sesterterpene 3-anhydro-6-epiophiobolin A (Duan *et al*., 2007).

Scytalone, a naphthalenone type PK, was present in all three strains and formed the largest cluster 1 in the FBMN (Figure 3A). This compound is a known intermediate in melanin biosynthesis and the presence of this compound concur with the presence of the BGC responsible for this metabolite in the genome data (Figure 3A). The cluster of nodes around scytalone (cluster 1) included ions with predicted molecular formulae and corresponding product ions that agreed with other polyketides. The other naphthalenone betaenone G was putatively annotated in this cluster as a constituent of Zt10 and IPO323. The BGC analysis (Figure 1C) indeed predicted the presence of a betaenone gene cluster in Zt10 and IPO323 strains, which thereby aligns with the metabolomics analysis. Betaenones A, B and C were however not annotated in the extracts. The polyketide cluster (1) included 27 ions shared by all three strains. In this cluster, only two ions seen as blue-only (*m/z* 323.3149 [M+H]^+^) and red-only (*m/z* 355.2904 [M+H]^+^) were specific to Zt10 and IPO323, respectively. We were not able to annotate these isolate-specific ions of polyketide origin, despite the use of multiple databases.

In the second largest cluster (cluster 2), 1,2-dilinoleoyl-SN-glycero-3-phosphocholine was annotated as belonging to a class of fatty acid (FA) compounds (Figure 3A) (Mills and Goldhaber, 2010). This MF had 23 ions with many putative positional isomers, which however could not be annotated (Table S10). Some sodiated ions were annotated as FAs in cluster 6 (Figure 3A) and were also observed in the FBMN.

Another class of compounds annotated in the *Z. tritici* metabolite network was the nonribosomal peptides of cluster 3. We annotated a class of diketopiperazines including cyclo Leu-Pro and cyclo Val-Leu (Figure 3A and Table S9) and the alloxazine lumichrome (Gao, et al., 2017) (Figure 3A). Cluster 5 was shared by all three *Z. tritici* isolates with 2 ions specific to Zt10. One of these ions could be annotated to be a sesterterpene 3-anhydro-6-epiophiobolin A (Figure 3A).

Overall, ∼ 30% of the detected ions could be annotated as known fungal metabolites, while the majority remains as putative new compounds to be described (Table S9). There was a negligible difference in the expression of compounds among the three field isolates (20/21 molecular families (or, the abbreviation of it: MFs) shared across the three *Z. tritici* isolates). Although Zt10 produced more ions (152 ions), IPO323 displayed more diverse chemistry and produced the only isolate-specific MF (cluster 13). This latter cluster comprises an ion, annotated to be a labdane diterpenoid sterebin Q4 (Figure 3A) and two un- annotated IPO323-specific singletons, *m/z* 373.2582 [M+H]^+^ and *m/z* 388.2692 [M+H]^+^, which may define a specific biochemical profile of the isolate. Taken together, our metabolomic study shows that the three isolates show comparable metabolites profiles in vitro and may harbor untapped molecules with biological functions.

### Removal of the histone modification H3K27me3 has little effect on BCG expression and metabolite production

Previous studies have revealed a prominent role of histone modifications in the regulation of fungal virulence factors, including effectors and secondary metabolites (Connolly *et al*., 2013; Soyer *et al*., 2014). We used a *Z. tritici* mutant deficient in the tri-methylation of histone H3 lysine 27 (H3K27me3) (Möller *et al*., 2019) to address if this histone mark likewise plays a prominent role in regulation of the identified BGCs in the *Z. tritici* genome. To this end, we re-analysed available transcriptome data (Möller *et al*., 2019) to assess the extent of differential gene expression between wild type *Z. tritici* and the methyltransferase mutant IPO323Δ*kmt6*, and we compared the metabolome profiles of wild type and mutant.

As previously reported (Möller *et al*., 2019), the wild type *Z. tritici* and the *kmt6* mutant do not differ substantially in gene expression (381 genes were found to be differentially expressed whereas 10169 genes did not show any significant difference in expression between the wild type and the mutant) (Table S10). Amongst the differentially expressed genes, only six are associated with the BGCs that we describe in this study, including a predicted transporter and a biosynthetic “additional” gene. None of these differentially expressed genes were predicted to be core biosynthetic genes. These results support previous findings suggesting that H3K27me3 does not play a prominent role in the regulation of secondary metabolite biosynthesis is *Z. tritici* (Möller *et al*., 2019).

To further investigate the role of facultative heterochromatin in the regulation of secondary metabolite production in *Z. tritici*, we compared the metabolomic profile of the wild type isolate IPO323 to the mutant *Δkmt6*, impaired in tri-methylation of the H3 lysine 27. Based on the FBMN, we generated the same set of MFs as produced for the field isolates (Figure S3). The data was visualized in a present-absent heatmap of all the detected molecular ions and their corresponding retention times and molecular clusters (Figure 4A). Altogether, 152 ions were produced and organized into 18 MFs (clusters) with 39 singletons (Figures 4A and S3). Importantly, no strain-specific MF was produced by the two strains, which confirms the transcriptome data and also indicate that H3K27me3 plays a minor role in regulating specific secondary metabolites in this fungus. Notwithstanding, the *Δkmt6* mutant produced 144 ions with 20 specific ions compared to the wild type, which produced 132 ions with 8 specific ions (Figure 4B). The majority of the ions, 124 representing approximately 81% of ions produced, were common to both strains (Figure 4B).

**Figure 4.**
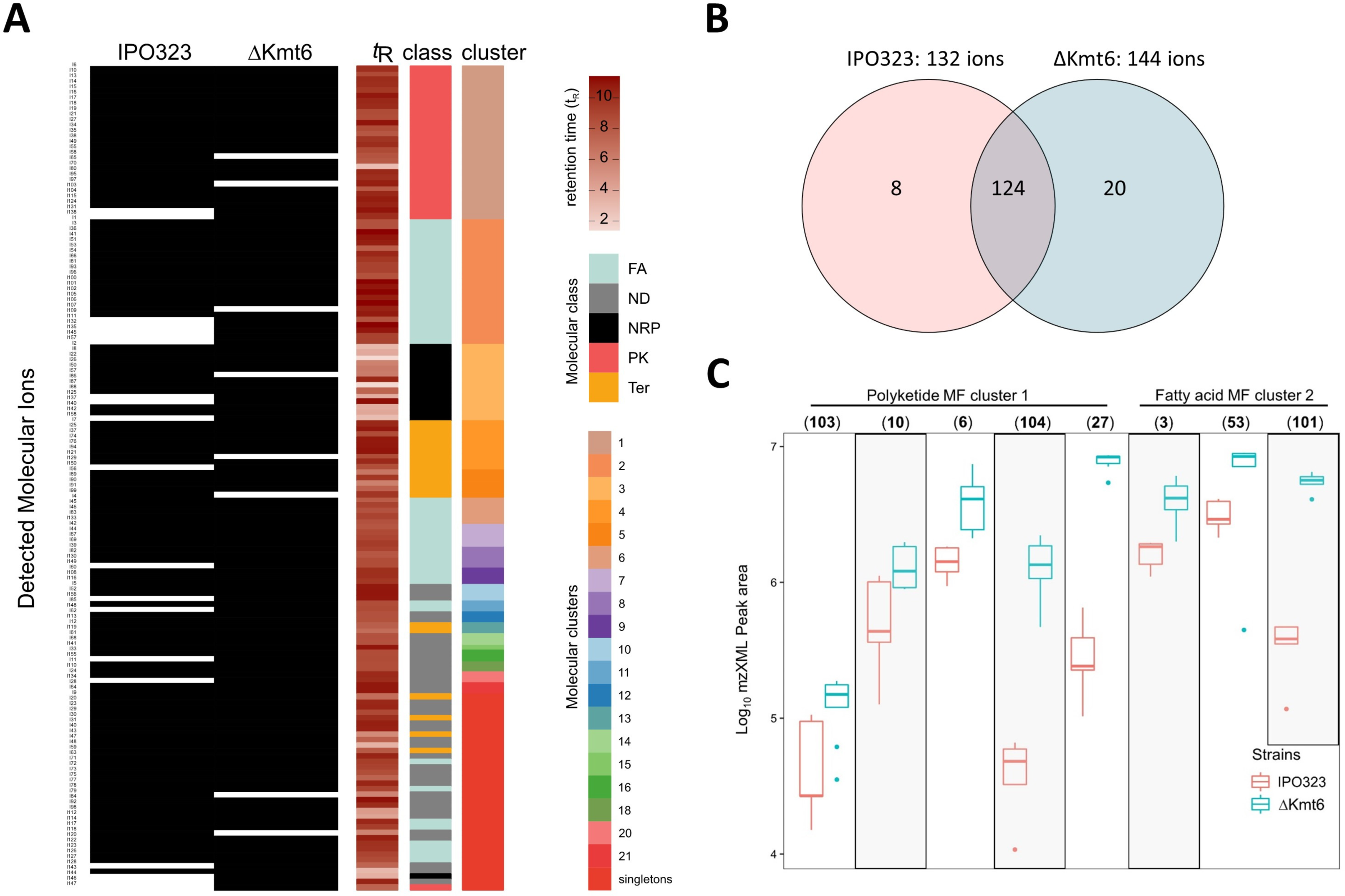
Comparative metabolomics of the wild strain IPO323 and ΔKmt6 mutant. **A)** Present-absent heat map from the UPLC-MS analyses of IPO323 and ΔKmt6 organic extracts showing the distribution of *m/z*, retention time, chemical classes (FA-fatty acid, NRP-nonribosomal peptide, PK- polyketide, TER-terpene and ND-not determined) and molecular family clusters. **B)** Venn diagram displaying specific and shared parent ions detected in the culture extracts of the IPO323 and Δkmt6 mutant. **C)** Box plots depicting the relative abundance of distinct molecular ions, in PK MF cluster 1 and FA MF cluster 2, between IPO323 and ΔKmt6 mutant of *Z. tritici*. Numbers in bracket represent unique ion IDs (as shown in Table S8 - (103) – *m/z* 395.2790 [M+H]^+^, (10) – *m/z* 321.2422 [M+H]^+^, (6) – *m/z* 379.3354 [M+H]^+^, (104) – *m/z* 397.2959 [M+H]^+^, (27) – *m/z* 323.2582 [M+H]^+^, (53) – *m/z* 518.3245 [M+H]^+^, (3) – *m/z* 520.3400 [M+H]^+^, (101) – *m/z* 784.5837 [M+H]^+^)).

Application of FBMN leveraged the MZmine feature detection and alignment tool to allow for relative quantification of peak ions produced by the IPO323 and *Δkmt6* mutant. Table S9 summarises the detected ions, their unique IDs and their relative abundance (peak areas) to assist comprehensive comparison of production titres of the major metabolites by the *Z. tritici* wild type and *Δkmt6*. Peak area box plots were generated for distinct ions detected in clusters 1 (polyketides) and 2 (fatty acids) to ascertain the effect of H3K27me3 on increasing the production titres of some compounds in the mutant strain. Several molecular ions from the polyketide MF cluster 1 showed an increased abundance in the mutant strain ΔKmt6 (Figure 4C, Table S9). This included 103 *m/z* 395.2790 [M+H]^+^, 10 *m/z* 321.2422 [M+H]^+^, 6b *m/z* 379.3354 [M+H]^+^, 104 *m/z* 397.2959 [M+H]^+^ and 27 *m/z* 323.2582 [M+H]^+^. In the fatty acid cluster 2, molecular ions 3 *m/z* 520.3400 [M+H]^+^, 53 *m/z* 518.3245 [M+H]^+^ and 101 *m/z* 784.5837 [M+H]^+^, were also expressed in relatively higher quantities in the *Δkmt6* mutant compared to the IPO323 wild type (Figure 4C). Overall, about 95% of the detected molecular ions were shared by the two strains with very minimal differences in terms of genes expressed under the specified cultivation conditions. In summary, while H3K27me3 does not contribute to the expression of specific expression of gene clusters and the production of individual metabolites, it enhances the production of several compounds quantitatively in the polyketide and fatty acid derived MFs.

## Discussion

The fungal species *Z. tritici* is one of the most devastating pathogens of temperate-grown wheat and causes severe yield losses to farmers in addition to high costs of crop protection measures. A challenge to the development of sustainable control strategies is the high level of genetic variation in field populations of *Z. tritici* that enables the fungus to rapidly adapt to changes in its environment. Genome and transcriptome studies have demonstrated that infection of *Z. tritici* relies not on major virulence determinants, but on a large repertoire of effector proteins that are produced to facilitate host invasion. In addition to proteinaceous virulence determinants, fungal pathogens also produce secondary metabolites during host infection. We addressed here whether the secondary metabolite profiles of three field isolates of *Z. tritici* also show a high variability consistent with the high variation of effector gene content and expression of the same isolates (Haueisen *et al*., 2018). We designed a pipeline for *in-silico* prediction of genes in three high quality genome assemblies of *Z. tritici* isolates (Feurtey *et al*., 2020). Based on these complete or near complete chromosome assemblies, we found very similar number of BGCs among the three isolates. Our updated prediction in this study provided a slightly lower number of BGCs than previously reported for the reference isolate IPO323 (29 versus 32), but comparable to predicted numbers of BGCs in other Dothideomycete genomes (Ohm *et al*., 2012). The conserved composition of BGCs in three field isolates of *Z. tritici* suggest that this pathogen does not rely on diverse chemistry as e.g., pathogenic and endophytic lineages of the plant-associated fungus *Epichloë* (Schardl *et al*., 2013), but rather on a conserved repetoire of metabolites.

Further annotation identified putative functions of some *Z. tritici* BGCs. Two of the predicted clusters show similarity to BGCs identified in other fungi and known to be involved in the production of the phytotoxins ustiloxin and betaenone. We find that genes located in the ustiloxin cluster are up-regulated during host infection and we hypothesize that the biosynthesis product plays a role in host colonization. The effect of ustiloxin has been documented on cell cultures and rice seedlings and the compound shows a variety of biological activities, including antimitotic activity whereby it can inhibit microtubule assembly (Wang *et al*., 2017). The relevance of this compound in the *Z. tritici*-wheat interaction is so far not known. For the betaenone cluster, we show up-regulation of one gene, exclusively in the isolates IPO323 and we hypothesize that this compound is of less relevance during host infection. Further, our analysis revealed that both fungal isolates harbor BGCs that encode for the synthesis of melanin. Melanin is encoded by broadly diverse fungal species and have been shown to have a critical role in fungal survival under stress conditions (Singaravelan et al 2008, Krishnan et al 2018) and could alter fungal pathogenicity (Chumley et al 1990, Liu and Nizet 2009). However, melanin-deficient strains of *Z. tritici* were not altered in their virulence or host colonization under test conditions (Derbyshire et al 2018), suggesting that melanin in *Z. tritici* may play other critical roles that require further testing.

We also show the up-regulation of genes in BGCs predicted to synthesize the antimicrobial and antifungal compounds gliotoxin, fusaridione and echinocandin. Recent studies have drawn the attention to the interaction of plant pathogens with their host microbiota (e.g. (Seybold *et al*., 2020; Snelders *et al*., 2020)). Fungal plant pathogens may produce a multitude of compounds to compete and exclude co- existing microorganisms in their host tissues. A recent study in the wilt pathogen *Verticilium dahliae* found an effector protein secreted by the fungus to manipulate host microbiota composition (Snelders *et al*., 2020). We predict that *Z. tritici* likewise produce antimicrobial and antifungal compounds to reduce growth of competing bacteria and fungi.

The gene cluster BGF_23 is predicted to be responsible for the synthesis of abscisic acid (ABA). Genes in this cluster show differential regulation across infection stages and are significantly up-regulated in all three isolates during the transition from biotrophic to necrotrophic growth. ABA is primarily known as a plant hormone playing a role in plant development and response to abiotic stresses. However ABA also has an antagonistic effect on jasmonic acid (JA) signaling and thereby reduce defense signaling (Anderson *et al*., 2004). Exogenous ABA was found to confer a strong increase in susceptibility of tomato seedlings towards the pathogen *Botrytis cinerea* (Audenaert *et al*., 2002). In the rice pathogen *Magnaporthe oryzae*, ABA was found to be a necessary component of the fungus virulence (Spence *et al*., 2015) and ABA has been qualified as an effector in both plants and animal pathogens (Lievens *et al*., 2017). It is worth mentioning that *Leptosphaeria masculans* produce ABA and the deletion of two genes involved in the production of ABA in this fungus did not alter its pathogenicity on canola. We speculate that biosynthesis of ABA during host colonization reduce JA-induced defense signaling and thereby facilitate pathogen colonization in the wheat - *Z. tritici* pathosystem as well. Functional characterization of the ABA core gene BGF_20:OG_1946 as well as an experiment with exogenous application of ABA will allow further testing of this hypothesis.

We applied metabolomics to characterize the biochemical profile of the three field isolates when propagated axenically. Consistent with the *in-silico* prediction, we observed highly similar metabolome profiles in the three field isolates (Figure 3B). Molecular networking revealed a high number of MFs common to all isolates. Although the overall metabolome profiles agree with the presence of BGCs predicted to produce these metabolites, there were other pathways for which the expected compounds were not detected. It is a well-known phenomenon that a majority of fungal BGCs are silent during cultivation in an artificial medium in the laboratory (Mao *et al*., 2018). Production and detection of the secondary metabolites rely (i) on its expression which may be influenced by the culture conditions, (ii) extraction techniques and (iii) production levels of the compounds which must be above the limit of detection for mass spectrometry or threshold employed for the automated feature detection. The lack of detection and annotation of ABA in the *in-vitro* extracts is thus consistent with the low expression of genes in the ABA cluster outside the plant (Duncan *et al*., 2015). We speculate that certain host cues induce the expression of the ABA BGCs during host infection. Interestingly, our metabolomics study also revealed some putative metabolites whose BGCs were not characterized in the genome analysis. The known phytotoxins such as trycycloalternarenes and ophiobolins, previously isolated from Dothideomycetes (Sviridov *et al*., 1991; Sugawara *et al*., 1998), were putatively annotated although BGCs encoding these compounds were not annotated in the genome data found. We speculate that this discrepancy is explained by lack of information in the databases that have formed the basis for our annotation analyses.

In some fungal species heterochromatin plays an important role in the regulation of secondary metabolites. In *Aspergillus nidulans*, loss of function of *CclA* a component of the COMPASS complex which methylates lysine 4 on histone H3, activated two pathways to produce the anthraquinone cluster of compounds-monodictyphenone and emodin, and the polyketides F9775A and F9775B (Bok *et al*., 2009). Here we included the mutant, IPO323Δ*kmt6* that is impaired in the tri-methylation of H3 lysine 27. While H3K27me3 is a global regulator of secondary metabolite production in *F. graminearum*, it appears to play a minor role in the regulation of BGCs in *Z. tritici*. This conveys with our previous RNAseq analyses of the same mutant showing a minor difference in transcriptional regulation (Möller *et al*., 2019). Rather, this histone modification plays a central role in the stability of accessory chromosomes in *Z. tritici* (Möller *et al*., 2019). It has been observed that in addition to awakening BGCs which are otherwise silent (Bok *et al*., 2009; Cichewicz, 2010), histone modification can impact the quantitative production of constitutive secondary metabolites. For example, the loss of *CclA* gene in *Aspergillus oryzae* enhanced the overproduction of the sesquiterpenes astellolides (Shinohara *et al*., 2016). The plant endophytic fungus *Pestalotiopsis fici CclA* mutant also increased the production of pestaloficiols and macrodiolide polyketides (Wu *et al*., 2016) providing further evidence for the relevance of histone modifications in secondary metabolite regulation in many fungal species. In *Z. tritici*, secondary metabolites are produced in the histone mutant, however we observe quantitative difference in the relative abundance of some metabolites whereby some metabolites are produced in higher quantities in the IPO323Δ*kmt6* mutant (Figure 4C). This finding suggests that some regulation occurs either directly or indirectly via H3K27me3 although not a main regulatory mechanism.

In conclusion, we find that the genome of the wheat fungal pathogen *Z. tritici* encode biosynthetic gene clusters that may produce a yet untapped diversity of metabolites. Our study demonstrates that the three *Z. tritici* isolates encode similar repertoires of BGCs that contrast the high variability in gene expression patterns, notably of effector genes and morphological phenotypes. Further, the in depth analysis of the transcriptional profiles of the annotated BGCs, as well as metabolomic profiles, did not reveal striking differences between the three field isolates, indicating that the three *Z. tritici* isolates share a common infection program. In contrast to *F. graminearum*, the histone methylation mediated by the histone methyltransferase Kmt6 plays a minor role in regulating the expression BGCs and production of secondary metabolites. This study reveal that *Z. tritici* encode for BGCs that potentially interfere with the plant immunity and compete against host-associated microbiome, Understanding at which infection stage these metabolites are produces will help to further decipher their function *in planta* and better understand the infection process of this pathogen.

## Experimental Procedures

### Genome data and BGC identification

To predict biosynthetic gene clusters (BGCs), the fungal genomes were submitted to antiSMASH fungi 4.0.2 (Blin *et al*., 2017). The Jaccard similarity network of BGC families was constructed using BiG-SCAPE (v1.0.0, cut-off distance 0.3) by mixing all classes (all versus all, (Navarro-Muñoz *et al*., 2020)). The network was plotted using R function ggnet2 from GGally package (https://ggobi.github.io/ggally). AntiSMASH analysis output could be accessed at https://zenodo.org/record/4592481. Gene function prediction were obtained from the data published in (Feurtey *et al*., 2020).

### RNAseq analyses

All transcriptomic datasets used in this study were previously published in (Haueisen *et al*., 2018; Möller *et al*., 2019). We re-analyzed the raw data with a focus using current annotation of BGCs. Adapter removal and read trimming were done with Trimmomatic v0.38 with parameters LEADING: 30 TRAILING:30 MINLEN:30 (Bolger *et al*., 2014). The alignment of the trimmed reads to each genome assembly (IPO323, Zt05 and Zt10) was done with HISAT2 version 2.2.1 with intron length set between 20 and 15000 (Kim *et al*., 2019) and the read counting with HTseq v0.11.2, both steps set with the option for reverse strand orientation (Anders *et al*., 2015). Genes annotated previously (Feurtey *et al*., 2020) as being reverse transcriptase or transposons were filtered out. Genes were defined as “not detected” when no reads aligned to them. Differential expression analyses were realized with the DESeq2 R package (Love *et al*., 2014). Thresholds for defining significance of differential expression were adjasted p-value <= 0.001 and log fold change > 2. The normalized expression was computed as Transcripts Per Million (TPM) from the read counts based on the exonic gene length and averaged over replicates for plots used in figures here. All scripts for the alignments, the TPM, the differential expression and the plots generation for the RNAseq analyses can be found in the supplementary file 1.

### Fungal extraction

In this study, we used three *Z. tritici* isolates that were previously described in (Haueisen *et al*., 2018). All fungal isolates were grown on solid YMS-agar medium (yeast extract 0.4% [w/v], malt extract 0.4% [w/v] and sucrose 0.4% [w/v] supplemented with 2% [w/v] bactoagar). The fungal isolates were incubated at 18°C for 15 consecutive days prior to extractions.

We pooled four solid cultures of *Z. tritici* and extracted metabolites with 400 mL EtOAc (PESTINORM, VWR Chemicals, Leuven Belgium) after homogenizing by an Ultra Turrax at 19000 rpm. Each EtOAc extract was washed twice with 200 mL of Milli-Q® (Arium®, Sartorius) water in a separatory funnel to remove salts. The EtOAc layer was then evaporated to dryness by a rotary evaporator (150 rpm at 40 °C). Dried extracts were solubilized in MeOH and filtered (0.2 μm filter) into storage vials and dried in vacuo. Each fungal strain was extracted in triplicate (biological replicates).

### UPLC-QToF-MS analysis

Chromatograms were acquired on an Acquity UPLC I-Class System coupled to a Xevo G2-XS QToF Mass Spectrometer (Waters®, Milford, MA, USA). Fungal extracts at concentrations of 1 mg/mL, were chromatographed on an Acquity UPLC HSS T3 column (High Strength Silica C18, 1.8 µm, 100 × 2.1 mm, Waters®) at 40 °C at a flow rate of 0.6 mL/min and injection volume of 0.5 µL in triplicates. A dual-solvent system used comprised of a mobile phase A: 99.9% MilliQ®-water / 0.1% formic acid (ULC/MS grade) and phase B: 99.9% acetonitrile (MeCN, ULC/MS grade, Biosolve BV, Dieuze, France) / 0.1% formic acid at a flow rate of 0.6 mL/min. The gradient was held at 1% B over 0.5 min, increased to 99% B over 11 mins, held at 99% for 3 mins, back to the starting condition over 0.5 min and maintained for 2.5 minutes.

MS^1^ and MS^2^ spectra were acquired in a data dependent analysis (DDA) mode with an electrospray ionization source in the positive and negative modes using the following parameters: A mass range of *m/z* 50–1200 Da, capillary voltage of 0.8 KV (1 KV in negative polarity), cone and desolvation gas flow of 50 and 1200 L/h, respectively, source temperature at 150 °C and desolvation temperature at 550 °C with sampling cone and source offset at 40 and 80, respectively. Collision energy (CE) was ramped: Low CE from 6 – 60 eV to high CE of 9 – 80 eV. As controls, solvent (MeOH) and EtOAc extract of sterile culture medium were injected under the same conditions. Only the data obtained from the positive mode were further analysed because the chromatograms revealed more complex profiles than in the negative mode.

### Molecular networking analysis of *Z. tritici* metabolites

UPLC-MS/MS data were converted to mzXML format using Proteo Wizard msconvert (version 3.0.10051, Vanderbilt University, Nashville, TN, USA) (Chambers *et al*., 2012) and then processed with MZMINE 2.33 [11,(Katajamaa *et al*., 2006; Pluskal *et al*., 2010). The mass detection was set to a noise level of 1000 for the MS1 level and 50 for MS^2^ levels. The chromatogram was built with ions showing a minimum time span of 0.1, minimum height of 3000 and *m/z* tolerance 0.01 (or 10 ppm). The chromatogram was deconvoluted with the baseline algorithm (minimum peak height 3000, peak duration 0.01 – 3, and baseline level 1000). Deisotoping of the chromatogram was achieved by the isotope peak grouper algorithm with *m/z* tolerance of 0 (or 10 ppm) and RT tolerance 0.2 min. All samples were combined in a peak list using the join aligner algorithm; the data were duplicate peak filtered and ions detected in the solvent and culture medium blanks were removed from the mass list. Only data with MS^2^ scans and within a retention time range (0 -11.5 min) were exported as csv (feature quantitative table) and mgf files and uploaded to the GNPS (Wang, et al., 2016) platform for feature-based molecular networking analysis (Nothias *et al*., 2020). The preprocessed data was filtered by removing all MS^2^ fragment ions within +/- 17 Da of the precursor *m/z* value. MS^2^ spectra were filtered by choosing only the top 6 fragment ions in the +/- 50 Da window throughout the spectrum. The mass tolerances were set to 0.05 Da for both precursor ion and fragment ion. A molecular network was then created where edges were filtered to have a cosine score above 0.7 and more than 6 matched peaks. Finally, the maximum size of a molecular family was set to 100, and the lowest scoring edges were removed from molecular families until the molecular family size was below this threshold. The spectra in the network were then searched against GNPS spectral libraries [14,(Horai *et al*., 2010; Wang *et al*., 2016). The library spectra were filtered in the same manner as the input data. All matches kept between network spectra and library spectra were required to have a score above 0.6 and at least 6 matched peaks. Cytoscape data was exported to Cytoscape version 3.6.14 (Shannon *et al*., 2003) to visualize the networks with nodes representing peak ions and edges representing similarities between peak ions. Molecular formula predictions were done with MassLynx version 4.1. for annotation of parent ions. Predicted molecular formulae were searched against databases such as Dictionary of Natural Product (http://dnp.chemnetbase.com), NP Atlas (https://www.npatlas.org) and SciFinder (https://scifinder.cas.org). Dereplicated peak ions were further checked by comparing the experimental fragments to in-silico fragments generated from the CFM-ID web server (Allen *et al*., 2014). The molecular networking job on GNPS can be found at https://gnps.ucsd.edu/ProteoSAFe/status.jsp?task=36c108371b1f41439b102a8ab75f1fce

## Acknowledgements

This research was supported by a personal grant from the State of Schleswig-Holstein to EHS. EHS furthermore acknowledges support from CIFAR.

## Supplementary Information

**Supplementary file 1:** Bioinformatic pipeline of the transcriptomic analyses

Table S1: Predicted BGCs from *Zymoseptoria tritici* isolate Zt05

Table S2: Predicted BGCs from *Zymoseptoria tritici* isolate IPO323

Table S3: Predicted BGCs from *Zymoseptoria tritici* isolate Zt10

Table S4: List of Biosynthetic gene family membership

Table S5: Summary statistics of RNAseq read mapping

Table S6: Differential gene expression *in planta*

Table S7: Transcripts per million read (TPM) *in planta*

Table S8: Detailed annotation of genes in the predicted ABA cluster

Table S9: Putative identification of metabolites in all *Zymoseptoria tritici* isolate

Table S10: Differential gene expression of IPO323 and kmt6 mutant during *in vitro* growth

Figure S1. **Expressed and non-detected genes during the course of infection.** Bar plot representing the number and percentages of expressed genes and genes with no detectable expression in the three strains of *Z. tritici*. Genes are separated in two categories: the genes predicted to belong to a BGC and all other genes. Numbers and proportions are given per infection stage.

Figure S2. **Transcriptomic profile of core biosynthetic genes during infection.** A dotted line represents change in expression level that is not significant whereas significant differences are represented by a full line. The line plots are labeled with the orthogroup number (written as: OGXXXX, X refers to a number) as well as with an identifier for the BGF (BGFXX, X refers to a number) to allow identification of core biosynthetic genes belonging the same cluster. In yellow are highlighted the BGF with some similarity to functionally characterized clusters of other species. The predicted products are as follow: BGF11 – Ustiloxin B; BGF17 – Echinocandin; BGF20 – Abscisic acid; BGF23 – Melanin; BGF28 – Fusaridione; BGF29 – Gliotoxin; BGF5 – Betaenone.

Figure S3. **Feature based molecular network (FBMN) of IPO323 and ΔKmt6 mutant**. FBMN generated from the Global Natural Product Social MN platform of crude organic extracts from IPO323 and Δkmt6 mutant of *Z. tritici*. Nodes represent molecular ions detected in the crude extracts with colour coding indicative of the relative abundance in each *Z. tritici* strain (red-IPO323 and green-Δkmt6 mutant). Some annotated compounds are displayed with their chemical classes: polyketide (PK), nonribosomal peptide (NRP), terpene (TER) and fatty acid (FA). Other annotations are displayed in Table S9.

